# The *Urtica Dioica* Agglutinin Prevents Rabies Virus Infection in a Muscle Explant Model

**DOI:** 10.1101/2021.05.21.441744

**Authors:** Xinyu Wang, Lisanne Terrie, Guanghui Wu, Els J.M. Van Damme, Anthony R Fooks, Ashley C Banyard, Lieven Thorrez, Dirk Jochmans, Johan Neyts

## Abstract

Infection with the rabies virus (RABV) results, once symptoms develop, in a 100% lethal neurological disease. Post-exposure prophylaxis (PEP), consists of a combination of vaccination and anti-rabies immunoglobulins (RIGs); it is 100% effective if administered early after infection. Because of the limited availability, alternatives for RIGs are needed. To that end we evaluated a panel of 33 different lectins for their effect on RABV infection in cell culture. Several lectins, with either mannose or GlcNAc specificity elicited anti-RABV activity of which the GlcNAc specific *Urtica dioica* Agglutinin (UDA) was selected for further studies. UDA was found to prevent entry of the virus into the host cell. To further assess the potential of UDA, a physiologically relevant RABV infection muscle explant model was developed. Strips of dissected swine skeletal muscle that were kept in culture medium could be productively infected with the RABV. When the infection of the muscle strips was carried out in the presence of UDA, RABV replication was completely prevented. We thus developed a physiologically relevant RABV muscle infection model. UDA (i) may serve as a reference for further studies and (ii) holds promise as a cheap and simple to produce alternative for RIGs in PEP.

## Introduction

Rabies virus (RABV), a member of the Lyssavirus genus, is a neurotropic virus that causes an acute and fatal encephalitis in humans and animals. The incubation period in most patients is 20-90 days [1], and, upon onset of symptoms, the disease is nearly 100% fatal. Post exposure prophylaxis (PEP), in the form of vaccination and injection of rabies immunoglobulins (RIG) in and around the wound shortly after exposure, can prevent infection [2]. However, RIGs are scarce in endemic regions [3,4] due to their short shelf-life, the need for a cold chain and the high cost of manufacturing. It is estimated that less than 2% of severely exposed patients (category III) receive RIGs worldwide [5,6].

Humans are most commonly exposed to rabies when bitten by rabid animals with infectious virus in their saliva. If the dermal barrier is breached, the virus infects muscle cells and gains access to neuronal cells through the neuromuscular junctions (NMJ). Once within neuronal cells the virus migrates from the peripheral nerves to the central nervous system (CNS) resulting in replication in the brain and the development of rabies. The disease is characterized by an altered mental status, hydrophobia, aerophobia, or inspiratory spasms [7]. To enable cell entry, the nicotinic acetylcholine receptor (nAChR) serves as one of the key receptor molecules for rabies virus binding and entry [8–10]. However, the nAChR tends to be concentrated at the postsynaptic muscle membrane in the NMJ, which may drive accumulation of rabies virions near the NMJ enhancing the likelihood of the virus entering the peripheral neuron or motor neuron [11,12]. To this end, it is important to inhibit RABV accumulation or replication in muscle cells and prevent infection of the nervous system. This mechanism of action is proposed for RIGs, which are injected in the tissue near the exposure wound.

Lectins are a group of carbohydrate-binding proteins from plants, fungi, animals or bacteria that can also be produced by recombinant DNA techniques [13]. Since the initial discovery of lectins more than 100 years ago, they became a focus of attention in many biological processes, including glycoprotein recognition, cell-cell communication, interaction with infectious agents, recruitment of leukocytes to inflammatory sites and tumor metastasis [13,14]. Several lectins, mostly with mannose and N-acetylglucosamine specificity elicit antiviral activity against HIV through binding to gp120 [15,16] and inhibition of viral entry [17]. HHA, a mannose specific lectin from *Hippeastrum Hybrid* has been reported to inhibit SARS-CoV at the stage of viral attachment and fusion [18]. UDA, a N-acetylglucosamine specific lectin from *Urtica dioica* inhibits SARS-CoV-2 infection of cells by binding to the spike protein [19]. Some lectins were shown to reduce infection of cells with the RABV [20,21].

We here identified several other lectins that inhibit RABV infection *in vitro*. To assess in a physiological relevant infection model, the potential of lectins as alternatives for RIGs, we established a RABV infection model in swine muscle strips. We demonstrate in this model that UDA, an N-acetylglucosamine specific lectin from the Stinging nettle (*Urtica dioica*) efficiently inhibits RABV infection in this muscle model.

## Materials and methods

### Cells, virus and lectins

Rabies virus SAD-B19-mCherry was obtained from the laboratory of Professor Anthony Fooks (Animal & Plant Health Agency, UK). BHK-21J cells (ATCC CCL-10™) were propagated in Dulbecco’s Modified Eagle Medium (DMEM) (ThermoFisher Scientific) with 10% fetal bovine serum (FBS) (Hyclone) and penicillin-streptomycin (P/S) (100 U/mL, ThermoFisher Scientific). When cells were infected with RABV, 2% DMEM was used. Lectins were obtained from Professor Els Van Damme (Ghent University, Belgium), then were dissolved in PBS at 1 mg/mL and stored at -20 °C.

### Antiviral assay

Compounds were added to cells in a 2-fold serial dilution from 100 μg/mL to 0.8 μg/mL, followed by virus at a multiplicity of infection (MOI) of 0.01 TCID_50_/cell. Antiviral activity was measured by quantification of the amount of mCherry fluorescence on day 5 post-infection (SPARK, Tecan, Belgium). Cell viability was determined by an MTS assay as described previously[22]. The half maximal effective concentration (EC_50_) of antiviral effect and half maximal cytotoxic concentration (CC_50_) was derived from the corresponding dose–response curves.

### Time of drug addition assay

UDA, at a final concentration of 25 μg/mL, was added to RABV infected (MOI =1) cell cultures. To determine if UDA has an impact on viral adsorption and attachment, cells were incubated with RABV at 4°C for 1h (−1 – 0 h) with or without UDA. After the incubation, to investigate if UDA inhibits viral entry, the unattached virus was washed away 3 times with PBS. Subsequently, cells were incubated at 37°C and UDA was added at different time points (0, 0.5, 1, 2, 4 h.p.i., Figure 2). Infected cells without compound treatment were defined as untreated controls, and untreated samples collected at 1 h.p.i. were considered as virus background. At 16 h.p.i., viral RNA was extracted from the cells (E.Z.N.A. Total RNA Kit I R6834-01, Omega BIO-TEK) and quantified by RT-qPCR analysis.

### Pre-incubation of RABV to lectins prior to infection

RABV (MOI=0.4) was incubated with UDA (25μg/mL) at 37 °C for 2 h. Then, the mixture of virus and compound was diluted and added to BHK cells and incubated at 37 °C (virus final MOI was 0.1, UDA final concentration was 1 μg/mL). After 3 days, the viral mCherry fluorescence was quantified (SPARK^®^, Tecan, Belgium). The result was defined as “Pre-Incubation Virus”.

### Pre-incubation of BHK to lectins prior to infection

The monolayer of BHK cells was pre-incubated with 25 μg/mL UDA at 37 °C for 2 hours. Then, the compound was diluted by adding the virus suspension and incubated at 37 °C (virus final MOI was 0.1, UDA final concentration was 1 μg/mL). After 3 days, the viral mCherry fluorescence was quantified as above. This result was defined as “Pre-Incubation Cells”

### Swine skeletal muscle explant culture and antiviral assay

The biceps femoris muscle was dissected from freshly euthanized pigs (3 months, female), obtained as residual tissue from ORSI Academy (Melle, Belgium). The muscle biopsy was dissected, parallel to the muscle fibers, in 2 cm long 2 mm thick strips using sterile scalpels. Tissue strips were maintained under tension using minutia pins (Entosphinx, stainless steel pins, diameter 0.25 mm) on sterile sylgard coated wells of a 6-well plate. During maintenance tissue strips were cultured (37°C, 5% CO_2_) submerged in DMEM with 10% FBS and P/S (100 U/mL), and in DMEM with 2% FBS and P/S during the antiviral experiments. For antiviral testing, muscle explants were incubated with 25 μg/mL UDA at 37 °C for 2h. Then they were inoculated with 2 × 10^4^ TCID_50_ RABV while the UDA concentration was kept constant at 25 μg/mL. After 4 h incubation, muscle explants were washed 5 times with PBS, and maintained with 25 μg/mL UDA for 6 days. For the untreated control, muscle strips were inoculated with same amount of virus but without UDA treatment. A sample of 300 μL of the culture SN was collected every day for untreated control for RT-qPCR analysis, and on day 6 for UDA treated sample for determination of infectious virus respectively.

### Muscle culture supernatant titration

The titer of infectious virus, present in swine muscle explant supernatant at day 6, was determined by 10-fold serial dilution on confluent BHK cells. On day 7, virus expressed fluorescence was quantified using microscopy and the TCID_50_ was calculated by the Spearman– Kärber method.

### Muscle explant viability assay

Muscle stripes were incubated with UDA at 25 μg/mL for 7 days. On each day, a resazurin-based toxicity assay was conducted: The medium of muscle stripes was removed and replaced by 1ml resazurin (PrestoBlue™ HS cell viability reagent P50200, ThermoFisher) working solution (10% PrestoBlue regent in PBS) followed by incubation at 37°C for 2h. Then, 200 μL resazurin working solution was transferred to a transparent 96-well plate. The fluorescence was measured at excitation 560nm and emission 590nm. In living cells, the cell-permeable, non-fluorescent resazurin is reduced to the red highly fluorescent resorufin. As resazurin is continuously converted into resorufin, the fluorescence is a quantitative indicator of the cell viability and cytotoxicity.

### Fluorescent immunostaining and microscopy of muscle

On day 6 p.i. swine skeletal muscle explants were washed 5 times with PBS, fixed in 4% formaldehyde for 1 h and stored at 4 °C in PBS until staining. Before imaging, they were permeabilized with ice-cold 80% acetone at room temperature for 1 h and washed 5 times in PBS. Next, muscle explants were incubated 1 h at 37 °C with FITC Anti-Rabies Monoclonal Globulin (800-092, Fujirebio) diluted 1:10 in PBS. After 5 times PBS washing, muscle explants were incubated in 5μg/mL Hoechst (H1399, Invitrogen) for 15 min at room temperature. Imaging was performed using the Leica DMi8 S Platform with Leica Application Suite X software (Leica, Heidelberg, Germany). All pictures were obtained with a 10× objective and z position between 1820 μm to 1950 μm.

### Real-time RT-qPCR

Rabies virus N gene was amplified by real-time quantitative PCR using iTaq™ Universal SYBR® Green One-Step Kit (BIO-RAD). The primers are as follow: 5’-TGGGCACAGTTGTCACTGCTTA-3’ (forward) and 5’-CTCCTGCCCTGGCTCAA-3’ (reverse). The standard curve was generated from the RNA extraction of the 1/10 diluted virus stock (2.2 × 10^6 TCID_50_/mL). A 20 μL qPCR reaction contains 4 μL extracted sample RNA or standard, 10 μL iTaq Universal SYBR® Green reaction mix, 0.25 μL reverse transcriptase and 600 nM of each forward and reverse primer. qPCR was performed in a Roche LightCycler 96 with the procedure: 10 min at 50 °C for reverse transcription, 1 min at 95 °C for polymerase activation and DNA denaturation, 40 cycles of 95 °C 15s and 62 °C 30s for PCR amplification. Viral copies were calculated based on a standard curve using control material.

### Statistics

Statistical analysis in this study was done by GraphPad Prism 9. A P-value less than 0.05 was considered as statistically significant. P values associated with each graph are indicated as: *, P value < 0.05; **, P value < 0.01; ***, P value < 0.001; ****, P value < 0.0001.

## Results

### Several lectins inhibit RABV in cell culture

The effect of 33 lectins on RABV infection of BHK cells was assessed using a reporter virus (SAD-B19-mCherry). Several lectins were identified that elicit antiviral activity (Table 1). The GlcNAc-specific agglutinin *Urtica dioica* Agglutinin (UDA) and the mannose-specific BanLec from *Musa acuminata* (banana) and PSA from *Pisum sativum* (pea) were most potent and selective. These activities were confirmed in a follow-up experiment with the same method as the screening (EC_50_s UDA: 4.6 μg/mL; BanLec: 6.0 μg/mL; PSA: 18 μg/mL) (CC_50_s UDA: 65 μg/mL; BanLec: 36 μg/mL; PSA: >100 μg/mL) (Figure 1). UDA (which we had available in larger quantities), was selected for further study. To explore at which step of the virus replication cycle UDA exerts its activity the lectin was added to the infected cultures at various times pre- and post-infection. When only present during the binding process at 4°C (from time -1 hr – 0 hr) (p.i.), no antiviral effect was measured. When UDA was however also present when the temperature was shifted to 37°C (which allows entry of the virus), the lectin prevented RABV infection. When addition of UDA to the cultures was delayed for one or two hours after the temperature shift, there was a gradual loss in antiviral activity (Figure 2). Together this indicates that UDA blocks RABV replication at the entry, but not the binding step. To investigate whether UDA interacts with the virus or the cells, either virus or cells were pre-incubated with 25 μg/ml UDA at 37°C for 2 hours after which the mix was diluted to a non-inhibitory concentration and was mixed with cells or virus, respectively. The RABV infection was quantified by measuring the viral fluorescence signal at 3 days p.i. Pre-incubation of the cells with UDA was more efficient in blocking the viral signal (*p* < 0.0001) than when virus was preincubated with UDA, indicating that the predominant mechanism of action may be the results of an interaction of the lectin with the host cell (Figure 3).

**Table 1.**
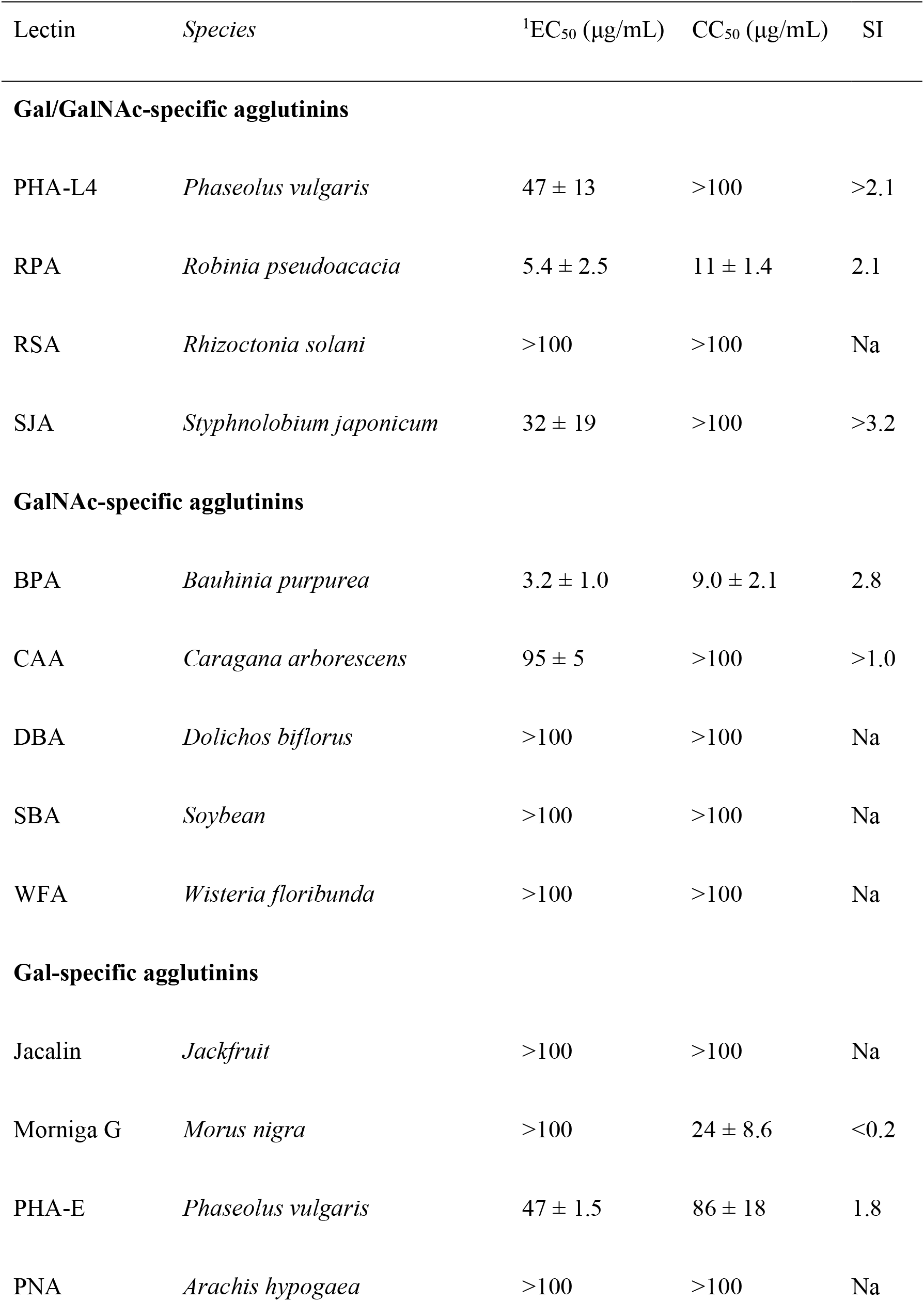
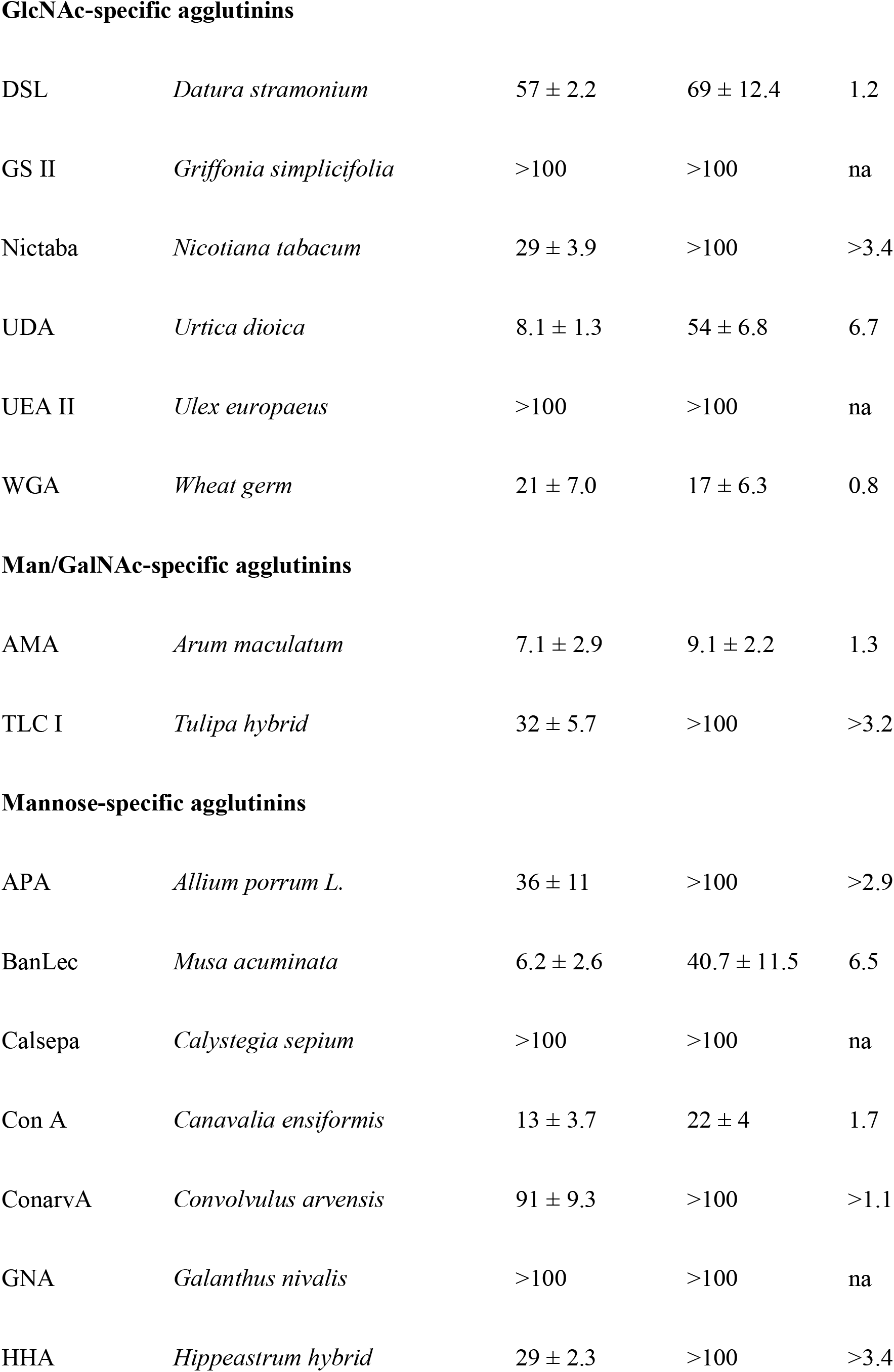
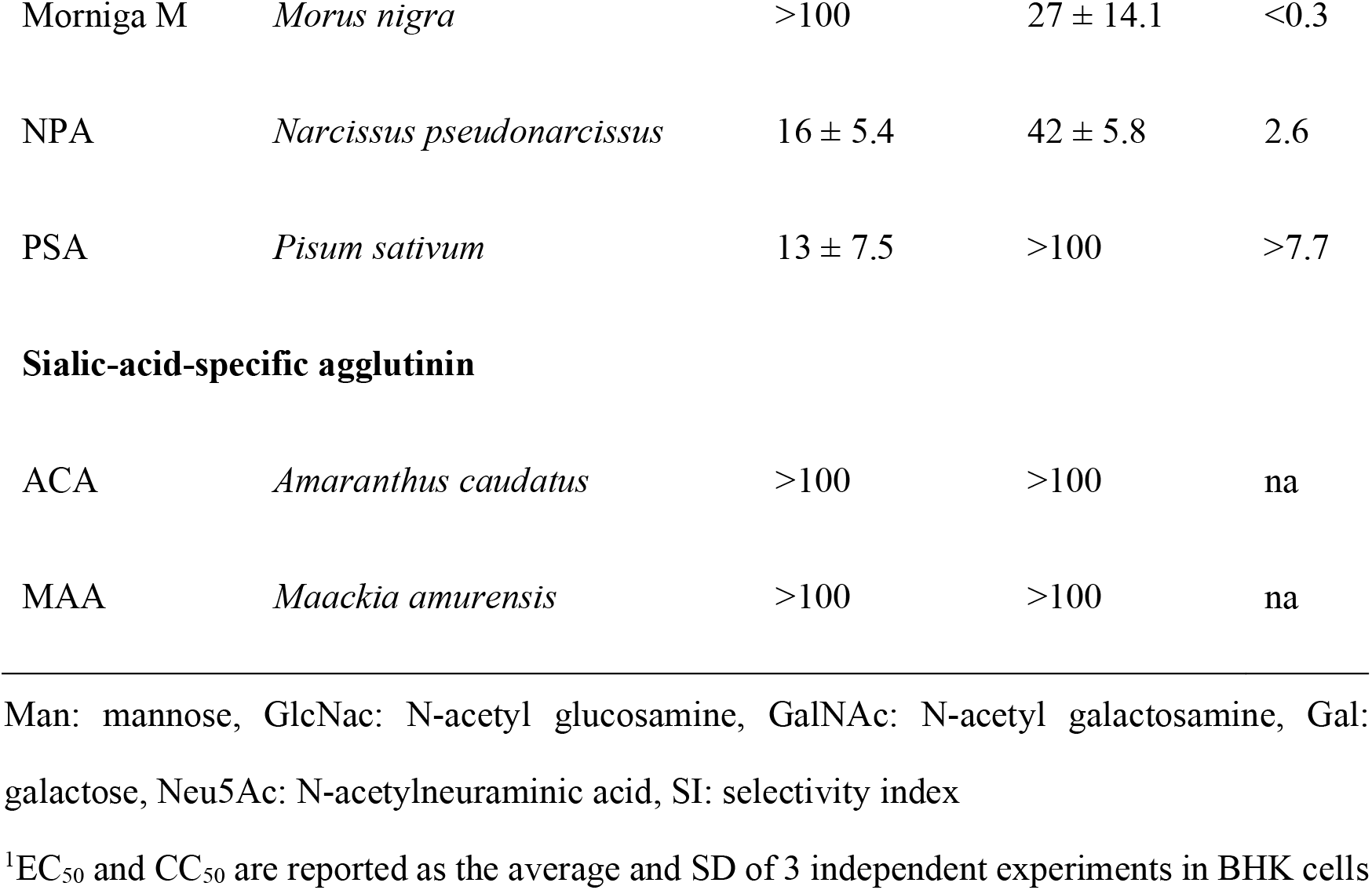
Antiviral activity of lectins against RABV.

| Lectin | Species | <sup>1</sup>EC<sub>50</sub> (µg/mL) | CC<sub>50</sub> (µg/mL) | SI |
| --- | --- | --- | --- | --- |
| **Gal/GalNAc-specific agglutinins** |  |  |  |  |
| PHA-L4 | *Phaseolus vulgaris* | 47 ± 13 | >100 | >2.1 |
| RPA | *Robinia pseudoacacia* | 5.4 ± 2.5 | 11 ± 1.4 | 2.1 |
| RSA | *Rhizoctonia solani* | >100 | >100 | Na |
| SJA | *Styphnolobium japonicum* | 32 ± 19 | >100 | >3.2 |
| **GalNAc-specific agglutinins** |  |  |  |  |
| BPA | *Bauhinia purpurea* | 3.2 ± 1.0 | 9.0 ± 2.1 | 2.8 |
| CAA | *Caragana arborescens* | 95 ± 5 | >100 | >1.0 |
| DBA | *Dolichos biflorus* | >100 | >100 | Na |
| SBA | *Soybean* | >100 | >100 | Na |
| WFA | *Wisteria floribunda* | >100 | >100 | Na |
| **Gal-specific agglutinins** |  |  |  |  |
| Jacalin | *Jackfruit* | >100 | >100 | Na |
| Morniga G | *Morus nigra* | >100 | 24 ± 8.6 | <0.2 |
| PHA-E | *Phaseolus vulgaris* | 47 ± 1.5 | 86 ± 18 | 1.8 |
| PNA | *Arachis hypogaea* | >100 | >100 | Na |
| **GlcNAc-specific agglutinins** |  |  |  |  |
| DSL | *Datura stramonium* | $57 \pm 2.2$ | $69 \pm 12.4$ | 1.2 |
| GS II | *Griffonia simplicifolia* | >100 | >100 | na |
| Nictaba | *Nicotiana tabacum* | $29 \pm 3.9$ | >100 | >3.4 |
| UDA | *Urtica dioica* | $8.1 \pm 1.3$ | $54 \pm 6.8$ | 6.7 |
| UEA II | *Ulex europaeus* | >100 | >100 | na |
| WGA | *Wheat germ* | $21 \pm 7.0$ | $17 \pm 6.3$ | 0.8 |
| **Man/GalNAc-specific agglutinins** |  |  |  |  |
| AMA | *Arum maculatum* | $7.1 \pm 2.9$ | $9.1 \pm 2.2$ | 1.3 |
| TLC I | *Tulipa hybrid* | $32 \pm 5.7$ | >100 | >3.2 |
| **Mannose-specific agglutinins** |  |  |  |  |
| APA | *Allium porrum L.* | $36 \pm 11$ | >100 | >2.9 |
| BanLec | *Musa acuminata* | $6.2 \pm 2.6$ | $40.7 \pm 11.5$ | 6.5 |
| Calsepa | *Calystegia sepium* | >100 | >100 | na |
| Con A | *Canavalia ensiformis* | $13 \pm 3.7$ | $22 \pm 4$ | 1.7 |
| ConarvA | *Convolvulus arvensis* | $91 \pm 9.3$ | >100 | >1.1 |
| GNA | *Galanthus nivalis* | >100 | >100 | na |
| HHA | *Hippeastrum hybrid* | $29 \pm 2.3$ | >100 | >3.4 |
| Morniga M | *Morus nigra* | >100 | 27 ± 14.1 | <0.3 |
| NPA | *Narcissus pseudonarcissus* | 16 ± 5.4 | 42 ± 5.8 | 2.6 |
| PSA | *Pisum sativum* | 13 ± 7.5 | >100 | >7.7 |
| **Sialic-acid-specific agglutinin** |  |  |  |  |
| ACA | *Amaranthus caudatus* | >100 | >100 | na |
| MAA | *Maackia amurensis* | >100 | >100 | na |

Man: mannose, GlcNac: N-acetyl glucosamine, GalNAc: N-acetyl galactosamine, Gal: galactose, Neu5Ac: N-acetylneuraminic acid, SI: selectivity index <sup>1</sup>EC<sub>50</sub> and CC<sub>50</sub> are reported as the average and SD of 3 independent experiments in BHK cells

**Figure 1.**
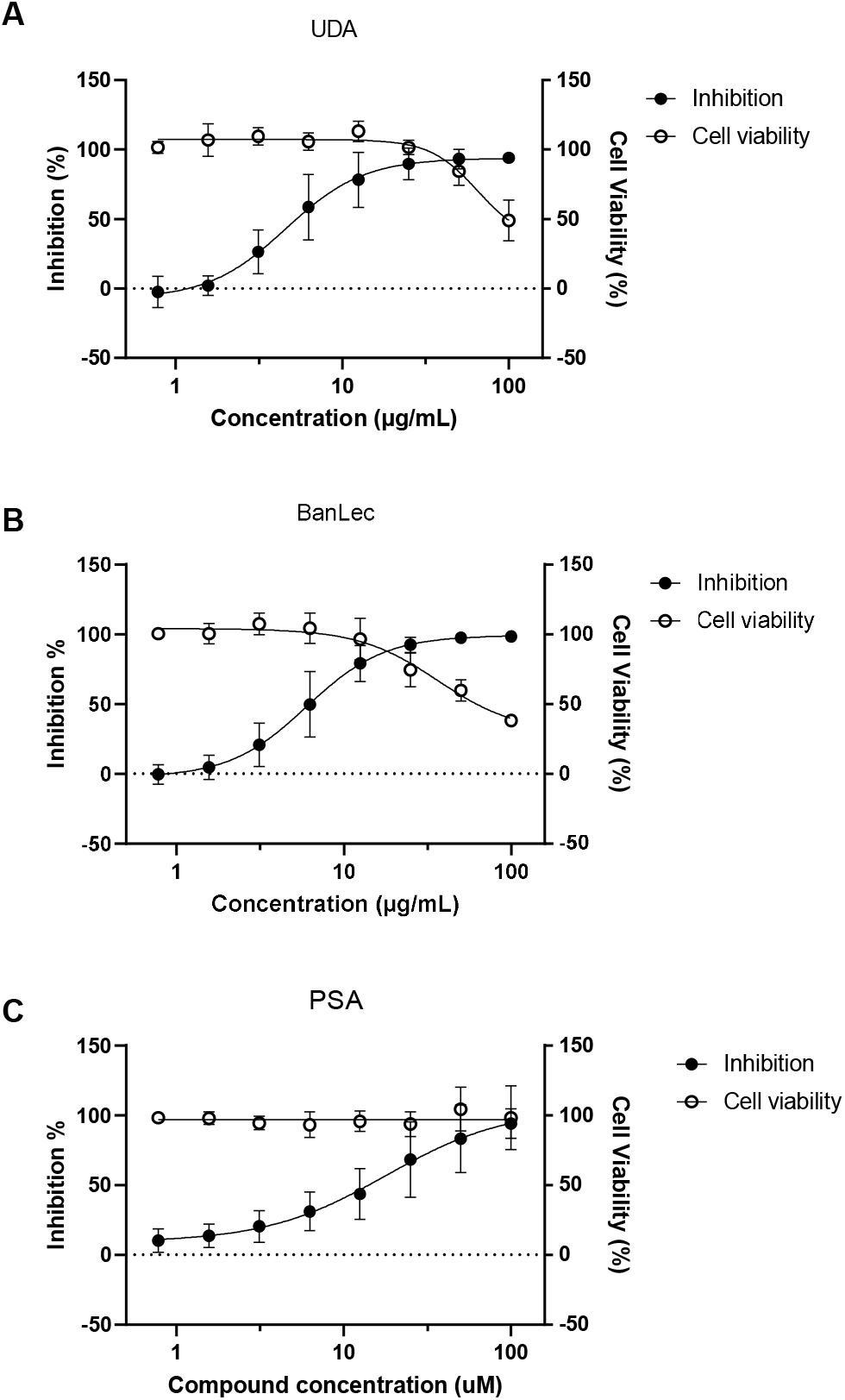
Anti-RABV activity of UDA, BanLec and PSA in BHK-21J cells. Serial dilutions of UDA (A), BanLec (B) and PSA (C) were added together with mCherry-RABV (MOI=0.01) to BHK cells. On day 5 p.i. the antiviral activity was determined by quantification of the virus induced mCherry fluorescence. In a parallel experiment the effects of the lectins on cell viability were determined using a viability staining (MTS). Averages and standard deviations of 3 independent experiments are presented. Fitting the dose-response curves indicates an EC_50_ of UDA of 4.6 μg/mL and a CC_50_ of 65 μg/mL; an EC_50_ of BanLec of 6 μg/mL and a CC_50_ of 36 μg/mL; an EC_50_ of PSA of 18 μg/mL and a CC_50_ of more than 100 μg/mL.

**Figure 2.**
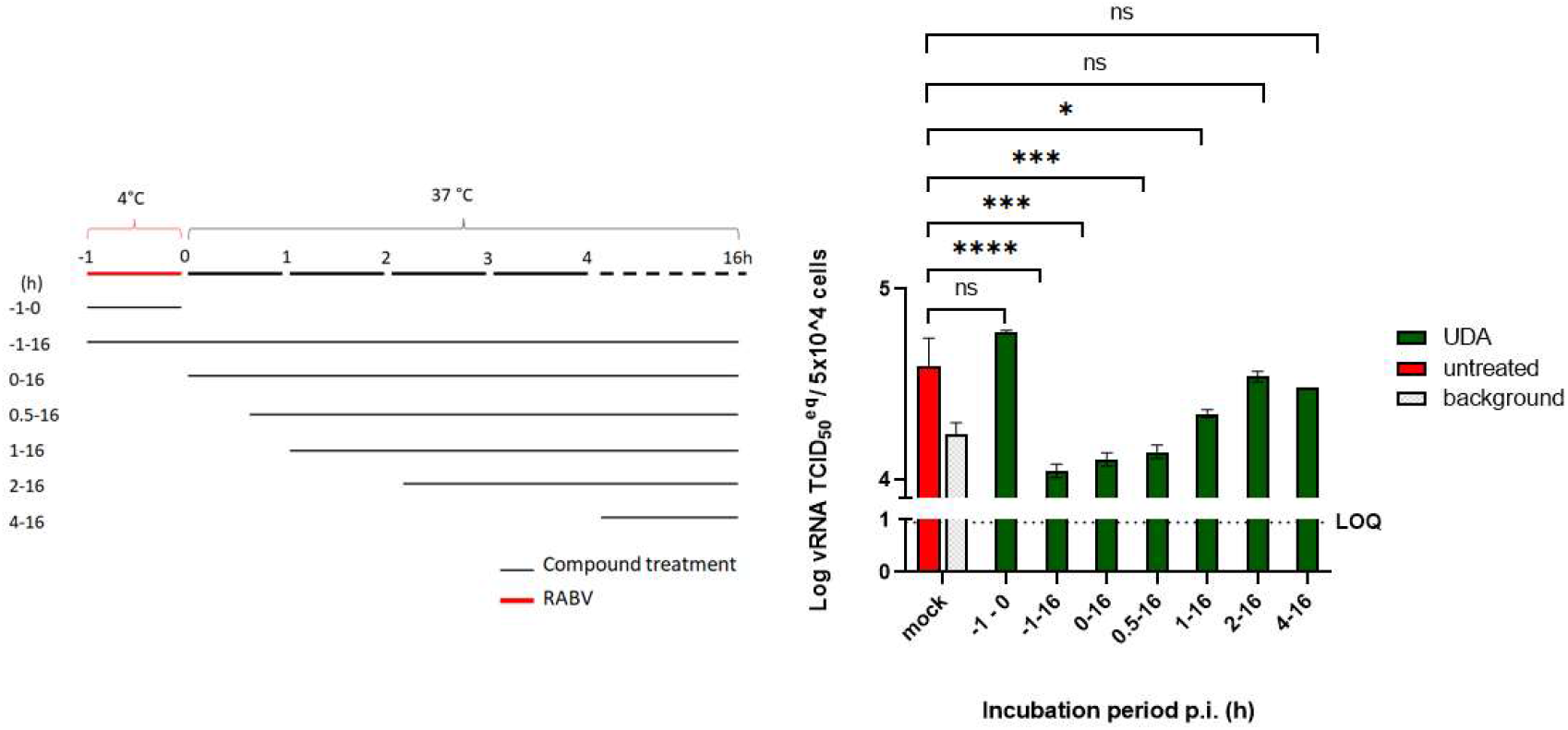
UDA blocks RABV entry. Time-of-drug-addition assay. BHK cells were incubated with RABV with or without UDA at 4°C for 1h (−1 – 0 h). After 1h, unattached virus was washed away and UDA was added to the cultures at different time points (0, 0.5, 1, 2, 4 hr p.i.). At 16 h.p.i., intracellular viral RNA was quantified by RT-qPCR. Infected-untreated samples collected at 1 h.p.i. were considered as the virus background. Each condition was tested in 3 independent assays and averages and STDEV are indicated. (*P < 0.05; *** P < 0.001, ****P < 0.0001)

**Figure 3.**
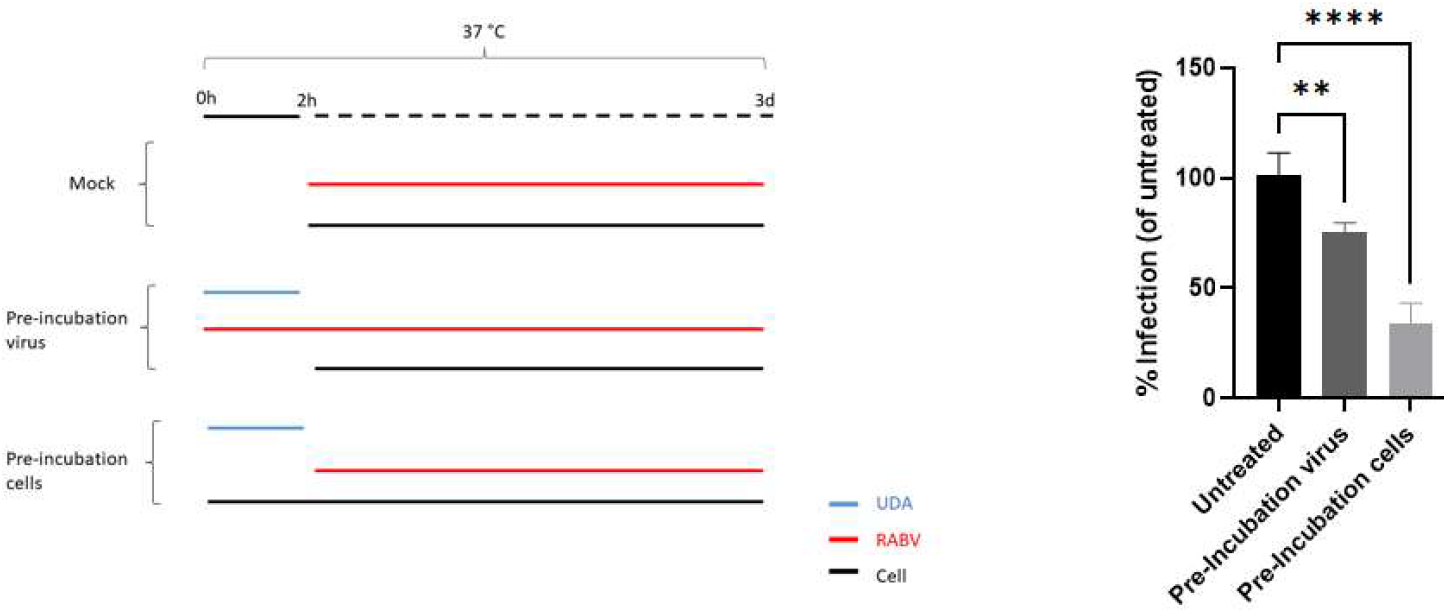
UDA inhibits RABV infection by mainly interacting with the cell. UDA at a final concentration of 25 μg/mL was pre-incubated with RABV or cells at 37°C for 2h. Next, UDA was diluted to a non-inhibitory concentration (1 μg/mL), and cells were further incubated with virus (MOI 0.1) for 3 days at 37°C. Virus infection (%) was determined relative to the untreated condition. Each condition was tested in at least 3 independent experiments and averages and STDEV were calculated. (** P < 0.01, ****P < 0.0001)

### RABV replicates in swine skeletal muscle explants

*Biceps femoris* muscle samples (2 cm long, 2 mm thick) were dissected from freshly euthanized 3-month-old female pig and maintained *ex vivo* under tension using pins (Figure 4A). In a first step we explored whether RABV can replicate in this muscle tissue. The explant cultures were inoculated with 2 × 10^4^ TCID_50_ of virus for 4h. Next, the viral inoculum was removed and the muscle strips were washed 5 times by PBS. The culture supernatant was collected every day for 6 consecutive days to determine the viral RNA load. On day 5-6 post-infection, viral RNA in the culture supernatant was >10 fold higher as compared to day 0 (Figure 4B). To confirm that infectious virus is produced, the culture medium obtained from the muscle cultures on day 6 was used to inoculate BHK cells. The mCherry red fluorescence expressed by the virus was observed in BHK cells infected with this day 6 inoculum (Figure 4C) demonstrating that swine skeletal muscles can be productively infected with RABV.

**Figure 4.**
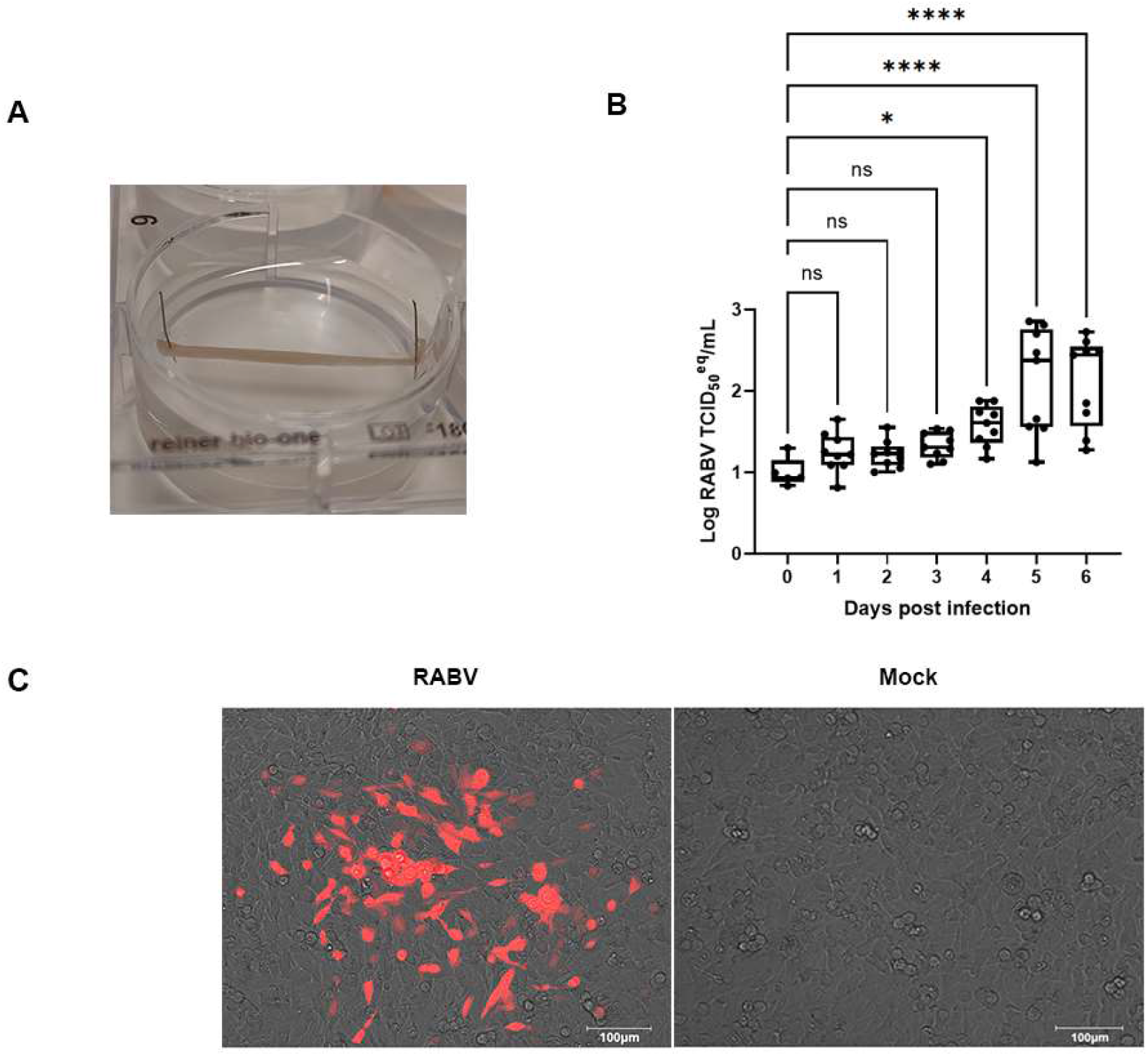
RABV replicates in swine skeletal muscle explants. (A) Picture of a swine muscle explant culture. The *biceps femoris* muscle was dissected from freshly euthanized pigs (3 months, female) and maintained under tension using pins. (B) Upon infection with RABV (SAD-B19-mCherry), culture supernatant was analyzed each day for levels of vRNA by RT-qPCR. Nine independent cultures were used, median and quartiles are indicated. (C) On day 6 p.i. the supernatant of infected (RABV) or non-infected (Mock) muscle cultures was transferred to BHK cells, and after 7 days of incubation, the fluorescence of the cells was visualized by microscopy (the picture is a representative culture of 3 independent experiments). Scale bar: 100 μm.

### UDA inhibits RABV infection in swine skeletal muscle

To investigate whether UDA can prevent RABV infection of muscle tissue, the muscle strips were pre-incubated with UDA (25 μg/mL) for 2 h followed by virus inoculation in presence or absence of UDA. After 4 h the virus inoculum was removed and new medium, with or without UDA (25 μg/mL), was added. On day 6 post-infection, the culture supernatant was collected for titration on BHK cells and the muscle tissues were fixed for immunofluorescence staining. UDA treatment resulted in a significant decrease in virus production in the medium (Figure 5B) and in antigen expression in the cells (Figure 5C). At a concentration of 25 μg/mL UDA had no adverse effect on the viability of the muscle strips during the 7 days exposure period (Figure 5A).

**Figure 5.**
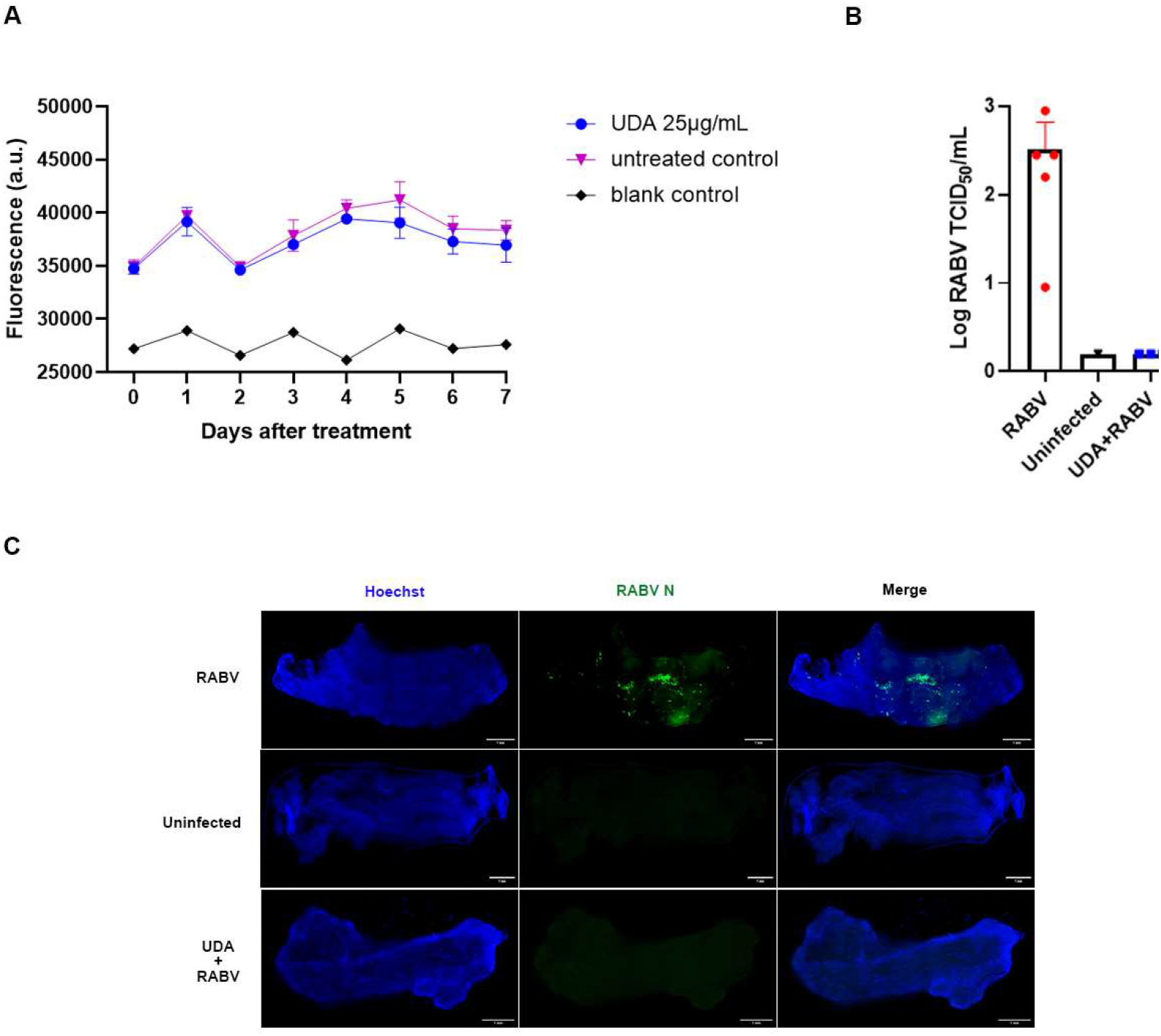
UDA inhibits RABV replication in swine skeletal muscle explants. (A) Assessing viability (using resazurin) of muscle explants that had been incubated for 7 days with UDA (25 μg/mL) at 37°C. The resazurin-based viability assay was conducted daily. Blank control: cultures without muscle strips. (B) Muscle explants were infected with RABV in the presence or absence of UDA (25 μg/mL). At 6 d.p.i. the culture supernatant was collected to determine infectious viral titers. (C) On day 6 p.i., muscle strips were fixed and RABV replication was visualized by staining for the viral N-protein (green), cell nuclei were stained with Hoechst (blue). Results of a representative example of 3 independent experiments is shown. Scale bar: 1mm.

## Discussion

Rabies immunoglobulins (RIG) are used, together with vaccination, in RABV post exposure prophylaxis (PEP) after high-risk exposure. RIGs provide passive protection by neutralizing the virus in the wound and surrounding tissues. Due to the high cost and supply shortages of RIGs, the WHO recommends since the 1990’s the development of alternatives [23]. Some monoclonals have been licensed for respectively the Indian (Rabishield and Twinrab) [24] and Chinese (Ormutivimab) market, but suffer obviously from the same shortcomings [5].

In an attempt to develop an alternative for RIGs, we explored whether a series of lectins (with various specificities), that are easy and cheap to produce and that do not require a cold-chain, may prevent entry of RABV in cells. We identified that the GlcNAc-specific agglutinin UDA from *Urtica dioica* (stinging nettle), the mannose specific lectins BanLec and PSA respectively from *Musa acuminata* (banana) and *Pisum sativum* (pea) as the most potent and selective inhibitors of infection within this series. UDA, (which was readily available to us) was selected for further studies. Others have reported that the mannose/glucose specific lectin Concanavalin A (con A) prevents RABV entry in cells, but we here observed that the antiviral and cytotoxic effects of con A are nearly overlapping [21]. Time-of-drug-addition studies and an experiment in which either the virus or the cells were preincubated with UDA indicate that this lectin prevents entry (not binding) of the virus in the cells and does so predominantly by interacting with the host cell. Lectins are characterized by their reversible binding to a specific mono- or oligosaccharide [25]. Several lectins have been reported to exert anti-bacterial and antiviral activities [13,14,26] and some lectins have been shown to elicit antiviral activity by binding with viral glycans of enveloped viruses [15–19] such as HIV and SARS-CoV2. Here, in contrast, we found that the mechanism by which UDA inhibits RABV infection stems primarily from an interaction with the host cells. Previous study shows that N-acetylglucosamine of the host cell is involved in an interaction of RABV with the cells [27]. UDA may thus interact with the cell in a way that prevents RABV entry but not binding.

Since RABV typically replicates in muscle tissue upon a bite from a rabid animal and before the virus enters the nervous system, we aimed to develop a physiologically relevant RABV infection model in muscle explants. We isolated intact swine skeletal muscles that were placed under tension in culture medium and that remained for at least 6 days metabolically active. When these muscles strips were infected with RABV, viral antigens were detectable in the tissue and infectious virus particles were released in the culture medium. This demonstrates that RABV infects and productively replicates in swine muscle explants. We then used this physiological relevant model to demonstrate that UDA can completely block RABV infection and subsequent replication in these muscle explants. Thus, the activity of UDA against RABV as observed in BHK cells also translates to muscle tissue demonstrating that this, and possibly, other lectins, may be further studied as alternatives for RIGs. Next, the efficacy of this, or ideally a lectin with more activity against RABV, or a combination of different lections, should be assessed in rodent models of RABV infection.

In conclusion, we developed a novel, physiologically relevant RABV infection model by using swine skeletal muscle explants and demonstrate that the plant lectin UDA can prevent RABV infection in this model. UDA and other lectins, may serve as a reference for further studies and may hold promise as an alternative for RIGs in PEP.

## Acknowledgements

We are grateful to Tina Van Buyten for excellent technical assistance.

## Author contributions

L. T. and L. T. isolated and dissected the muscle stripes from the freshly euthanized pigs. G. W. provided the rabies virus-mCherry. E. V. D. provided the library of lectins. A. R. F. and A. C. B. advised suggestions for writing. X. W., D. J. and J. N. designed the study and wrote the manuscript.

